# Molecular Identification and Antimicrobial Potential of Endophytic Fungi Against Some Grapevine Pathogens

**DOI:** 10.1101/2024.08.06.606939

**Authors:** Lava H. Nashat, Raed A. Haleem, Shayma H. Ali

**Affiliations:** Department of Biology, College of Sciences, University of Duhok, Duhok province, Kurdistan region-Iraq; Department of Plant Protection, College of Agricultura Engineering Sciences, University of Duhok, Duhok province, Kurdistan region-Iraq

**Keywords:** Biological agents, Grape. Phylogenetic analysis, Endophyte, ITS, SSU

## Abstract

In the Duhok province of the Kurdistan region, the plant species *Vitis vinifera* has been explored as a habitat for diverse endophytic microorganisms across various ecological environments. During the period from 2021 to 2022, a total of 600 samples were collected from four distinct locations: Bagera, Besfke, Barebhar, and Atrush. From these samples, twelve endophytic fungal species were isolated, including *Aspergillus flavus, Botryosphaeria dothidea, Fusarium oxysporum, Fusarium ruscicol, Fusarium venenatum, Chaetomium globosum, Clonostachys rosea, Mucor racemosus, Penicillium glabrum, Aspergillus terreus, Aspergillus ustus*, and *Aspergillus niger*. These fungi are recognized for their potential as biocontrol agents against grapevine trunk diseases and grape rotting fungi, which pose significant risks to grapevine health and productivity. Based on ITS and SSU sequencing, molecular identification confirmed these fungi’s presence with sequence identities ranging from 99% to 100%. Phylogenetic analysis revealed that these endophytes could be categorized into five main clusters (A, B, C, D, and E), showing high intra-group similarity. Utilizing the Dual Culture method, the endophyte *Paecilomyces maximus* demonstrated a 70.83% inhibition rate against *Ilyonectria destructans*. In the Food Poisoning method, *A. flavus* and *P. maximus* emerged as the most effective inhibitors of *Ilyonectria destructans*, whereas *A. terreus, M. racemosus*, and *P. maximus* achieved complete inhibition (100%) of *Botrytis cinerea*. Additionally, *M. racemosus* was identified as the most effective biocontrol agent against *Neoscytalidium dimidiatum*.

## 1. Introduction

Endophytic fungi engage in complex interactions with their host plants, often providing mutual benefits such as enhanced growth, stress tolerance, and disease resistance. In the context of grapevines, endophytic fungi are not only integral to plant health but also hold potential for drug discovery and improving plant resilience against various stresses like disease, drought, and salinity [1]. Recent studies have demonstrated that fungal endophytes can colonize plant tissues through the development of hyphae, establishing communities within specific ecological niches [2,3]. This colonization can result in various benefits for the host, including enhanced nitrogen fixation, protection against pathogens, and increased resilience to environmental stressors [4]. Grapevine cultivation is a significant agricultural practice worldwide, including in the Duhok Province of the Kurdistan Region. Most previous studies have focused primarily on the endophytic fungal diversity in cultivated grapevines, with less emphasis on wild grape species [5]. A comparative study of these grapevine types could yield valuable insights into the domestication process, host specificity, distribution patterns, and beneficial impacts of endophytes, as well as elucidate the diversity of endophytic fungal species.

In Duhok Province, grapevines face numerous biotic and abiotic challenges as in other vineyard regions. Adverse environmental conditions can significantly affect grapevine health and the quantity and quality of grape production [6]. The impact of cultivation practices on endophytic fungal populations in grapevines remains underexplored Furthermore, grapevines are susceptible to several fungal pathogens that cause significant crop losses. Historically, sodium arsenate was used to manage grapevine trunk diseases (GTDs), but its use was discontinued due to its severe toxicity to humans and the environment [7]. Consequently, there is a pressing need to explore alternative, safer methods for controlling GTDs. Recent research has highlighted the potential of endophytic fungi as biological control agents for effective disease management [8].

Previous research has documented the antagonistic effects of endophytic fungi on grapevine pathogens, including *Alternaria alternata, Aspergillus terrus, Botryosphaeria dothidea, Mucor racemusus, Paecilomyces*, and *Curvularia*. These fungi have shown promise in suppressing diseases caused by pathogens like *Cytalidium, Macrophomina, Neoscytalidium*, and *Botrytis* [5,9]. In Iraq, grapevine decline caused by *Cylindrocarpon destructans* was frequently observed and successfully controlled in vitro using two bioagents: *Trichoderma harzianum* and *Clonostachys rosea* [10,11]. Our study seeks to expand the ecological knowledge of grapevine endophytic fungi and explore their roles as biocontrol agents, providing a foundation for future research on sustainable disease management strategies. Advances in fungal identification have evolved from traditional morphological methods to modern molecular techniques, such as DNA sequencing, which offer more precise fungal classification.

## 2 Materials and methods

### 2.1 Sample collection and study site

Samples were collected from four distinct locations within Duhok city (Bagera and Besfke, Baribhar and Atrush) between 2021 and 2022. A total of 600 samples were obtained, including various plant parts: trunk (ST), nodes (N), internodes (IN), roots (R), leaves (L), petioles (P), bunch pedicels (BP), rachis and pedicel (RP), and mature fruit (MF). These samples were sourced from plants trained using two different systems: the Head training system and the Cordon training system. The collected samples were placed in sterile plastic bags and stored in a controlled environment inside a cool box. These samples were then handled in a sterile manner within 24 hours of collection and maintained them at a temperature of 4°C until use [12].

Surface disinfection is an essential step in the isolation of endophytic fungi, as it effectively removes epiphytic microorganisms that may impede the identification of true endophytes. The disinfection protocol involves several critical steps: initially, samples are washed with tap water to remove visible contaminants such as dust, debris, yeasts, and filamentous fungi [13]. Following this, the samples are submerged in 70% ethanol for 30 seconds, then treated with sodium hypochlorite (NaOCl) for 2 minutes, and subsequently submerged in 70% ethanol for 15 seconds. Finally, the samples undergo multiple rinses with sterilized, autoclaved distilled water. This comprehensive method ensures that the plant materials are thoroughly disinfected and prepared for further analysis.

### 2.2 Culturing of endophytic fungi

The plant samples were plated on Potato Dextrose Agar (PDA) supplemented with streptomycin (100 mg/L) and incubated at 25°C for 10 days. Fungal growth was monitored daily. The isolates were grouped based on their morphological characteristics. To facilitate further identification, the hyphal tips of fungal colonies were transferred to PDA slants [14]. This isolation method, as recommended by multiple sources [14,15] involves collecting hyphae from the periphery of the fungal colonies and introducing them into a fresh culture medium with antibiotics. The fungal isolates were classified based on their macro- and micromorphological characteristics following purification [16,17].

### 2.3 Molecular identification

#### 2.3.1 DNA extraction

Genomic DNA was extracted with the established CTAB methods [18]. Briefly, the whole endophytic fungus was isolated from pure fungal cultures. The Erlenmeyer flasks were used to introduce endophytic fungal mold strains into 200 ml of potato dextrose broth. The flasks were incubated for a period of ten days at a temperature of 25°C, cell walls of fungal mycelia were broken down by liquid nitrogen [19].The grinded fungi were subjected to CTAB protocol and purified using phenol:chloroform:isoamyl (25:24:1). Nucleic acids were precipitated by isopropanol. The precipitated nucleic acids dissolved in free nuclease water and stored at -4°C until used. The quantity and the quality of the extracted DNA were measured by a NanoDrop 2000UV-spectrophotometer, at two distinct wave length 260 nm and 280 nm. The purity of DNA was determined using the absorbance ratio A260\A280.

#### 2.3.2 PCR amplification of ITS and SSU regions

Amplification of ITS and SSU regions was carried out by PCR in thermos cycler machine (Eppendorf AG\USA) using ITS1 primer (5′-TCCGTAGGTGAACCTGCGG -3′) and ITS4 (5′-TCCGCTTTATTGATATGC-3′) [20]. PCR steps are an initial denaturation cycle for 5 minutes at 95°C, followed by 35 cycles starting with denaturation at 94°C for 1 minute, and annealing at 55°C for 45 sec. and extension at 72°C for 45 sec followed by a final extension for 7 minutes at 72°C. while the PCR conditions of NS1(5′-GTAGTCATATGCTTGTCTC-3′) and NS4(5′-CTTCCGTTCAATTCCTTTAAG-3′) [20], were as follows: initial denaturation cycle at 95 °C for 5 minutes followed by 40 cycles beginning with denaturation for 35 sec at 96°C, annealing at 55°C for 30 sec., extension for 1minute at 72°C followed by a final extension at 72°C for 5 minutes. The amplified PCR products of ITS (500bp) and SSU (1150bp) were separated by agarose gel electrophoresis, and observed under a UV-transilluminator.

#### 2.3.3 Sequencing of ITS and SSU regions

Qiagen Minielute purification kit were used to purify PCR products following the manufacturers instruction. The purified PCR products had sequenced by Macrogen Incorporation (Seoul, South Korea), using an ABI3730 XL automatic DNA analyzer and the primer pair ITS1, ITS4 and SSU (NS1, NS4).

#### 2.3.4 Analysis of ITS and SSU regions and species identification

Geneious, version R 8.1 Biomatters 14 and BioEdit version 7.2 software programs were applied to edit, analyze, trim, and verify the sequenced fragments of both forward and reverse primers for both regions and saved in fasta format. ITS and SSU amplicon sequenced fragments have been compared with available sequences at NCBI (National Center for Biotechnology Information). using the Basic Local Alignment Search Tool (BLAST) against ITS and SSU sequences of type isolates (www.ncbi.nlm.nih.gov/ BLAST, the nucleotide sequences showed ≥ 99 % similarities.

### 2.3.5 Phylogenetic analysis

BioEdit and MEGA software programs [21], have been used for phylogenetic analysis and nucleotide sequence alignment. The ClustalW algorithm using the default parameters has been used to align sequences. The Neighbor-joining method is applied to construct a phylogenetic tree.

### 2.4 In-vitro Antagonism potential of bioagents

#### 2.4.1 Dual Culture Method

Each set of endophytic and pathogenic fungal mycelium plugs was placed on a PDA plate, with a distance of 4 cm between them. The distance from the colonies’ margins to the plugs was measured as (0.5 cm). The endophytic fungus was positioned on one side of the petri dishes, 2 cm from the edge. Conversely, circular pieces (5 mm in diameter) were extracted from the outside edge of each mature pathogenic fungal culture. The discs were after that placed in an incubator at a temperature of 25±2ºC for 10 days. Three individual plates were created for each isolation. The control treatments consisted of pure fungal cultures. After the pathogen development in the control treatment was finished, the radial growth of all treatments was evaluated. The percentage of inhibition of mycelial growth of the test organisms compared to the control was calculated using the formula: I (%) = (C-T)/C×100, where I represents rate inhibition, C represents the rate of controlled growth and T represents the rate of growth in the treatment [22].

#### 2.4.2 Food poisoning method

The culture from each isolated bioagent was transferred and placed in a 250 mL Erlenmeyer flask containing 50 mL of potato dextrose broth (PDB). The flask was then incubated in the dark at a speed of 150 rpm for 72 hours. Following the incubation period, the fungal spores in each Bioagents culture were eliminated by filtration using Whatman N°4 filter paper and a 0.45 μm Millipore membrane, as per the modified methods of (Frighetto and Melo, 1995). Petri plates were filled with Potato Dextrose Agar (PDA) medium and 10 mL of each filtrate, each with a 50% concentration, was applied to the plates. After solidification, a 7 mm culture disc of pathogenic fungus was placed in the center of a Petri plate. The mycelial diameter of the pathogenic fungus was measured using two perpendicular orientations after the pathogens were inoculated in the experimental group’s petri plate. The rate of suppression of mycelial growth in the test organism compared to the control were calculated using the formula proposed by [22].

## 3. Results and discussion

### 3.1 Prevalence of endophytic fungi

Twelve endophytic fungi were identified from healthy *Vitis vinifera* plants in four regions of Duhok province (Bagera, Besfke, Baribhar, and Atrush). The following fungus has been isolated: *Aspergillus terreus, Aspergillus ustus, Aspergillus niger, Aspergillus flavipes, Botryosphaeria dothidea, Fusarium oxysporum, Fusarium ruscicol, Fusarium venenatum, Chaetomium globosum, Clonostachys rosea, Mucor racemosus*, and *Penicillium glabrum. Mucor racemosus* is the only fungus among them that is not classified under the Ascomycota group; which is belong to the Zygomycota. with the highest occurrence in Besfki, Baribhar and Atrush (100%),(83.3), and (83.3%) respectively, while *Clonostachys rosae* most occur in Bagera (100%) Table 1.

**Table 1.** The frequency of endophytic fungi in four distinct areas within the Duhok province.

| Endophytic Fungi/ Locations | Bagera | Besfke | Baribhar | Atrush |
| --- | --- | --- | --- | --- |
| *Aspergillus terreus* | 33 | 83.3 | 50 | - |
| *Aspergillus ustus* | - | 33 | 33 | - |
| *Aspergillus niger* | 66.7 | 83.3 | 66.7 | 66.7 |
| *Aspergillus flavipes* | 50 | 50 | 83.3 | - |
| *Botryosphaeria dothidea* | 33 | - | - | 66.7 |
| *Fusarium oxysporum* | 66.7 | 66.7 | 33 | 33 |
| *Fusarium ruscicol* | - | 33 | - | - |
| *Fusarium venenatum* | - | 50 | - | - |
| *Chaetomium globosum* | 33 | 33 | - | 33 |
| *Clonostachys rosea* | 100 | 16.7 | - | 50 |
| *Mucor racemosus* | 50 | 100 | 83.3 | 83.3 |
| *Penicillium glabrum* | 66.7 | 50 | 33 | 50 |

**Table 2.** Taxonomic units identified using molecular identifications:

| # | Isolated | Similarity | ITS<br>accession<br>number | SSU accession number |
| --- | --- | --- | --- | --- |
| 1 | *Fusarium oxysporum* | 99.81 | PP410452 | PP342393 |
| 2 | *A.terrus* | 99.82 | PP410453 | PP342394 |
| 3 | *A.ustus* | 100 | PP410454 | PP342395 |
| 4 | *Clonostachys rosea* | 100 | PP410455 | PP342396 |
| 5 | *Mucor racemosus* | 100 | PP410456 | PP342397 |
| 6 | *A.niger* | 100 | PP410457 | PP342398 |
| 7 | *A. flavipes* | 100 | PP410458 | PP342400 |
| 8 | *Chaetomium globosum* | 100 | PP410459 | PP342401 |
| 9 | *Penicillium glabrum* | 100 | PP410460 | PP342402 |
| 10 | *Fusarium ruscicola* | 100 | PP410461 | PP342403 |
| 11 | *Botryosphaeria dothidea* | 99.80 | PP410462 | PP342404 |
| 12 | *Fusarium venenatum* | 100 | PP410463 | PP342405 |

### 3.2 Molecular Identification and Phylogenetic Analysis

The qualification and quantification results showed the concentration of the isolates were ranged between (200-1300 ng/µl) with a purity between (1.6-1.82). The obtained sequenced results of ITS and SSU revealed a diversity of endophytic fungi from all parts of the grapevine, five fungal from leaves five from inter nods, three from nods, three from tendrils, four fungal from fruits, and six fungal isolated from roots.

Blast analysis of SSU and ITS regions resulted in 7 endophytic fungi genera Fig1. Different evolutionary groups shown in the phylogenetic relationships between different fungal species, as seen in the ITS and SSU genetic regions trees. Group A comprises *Aspergillus species* (A. *terreus, A. flavipes,A. niger*, and *A. nidulans*), which exhibit significant confidence in their connections with high bootstrap values indicating close relatedness. *Aspergillus species* and *Penicillium glabrum* in Group B have a tight but different relationship. Different from another fungus, *Botryosphaeria dothidea* is the representative of Group C. *Clonostachys rosea, Chaetomium globosum*, and many *Fusarium species* (*F. rusicola, F. oxysporum, F. venenatum*) comprise Group D. Fusarium forms a well-supported subgroup. The most distantly related species in Group E is *Mucor racemosus*, suggesting an early split from the common ancestor. This tree illustrates the diversity of fungal evolution by highlighting the closer relationships between *Aspergillus species*, the different evolutionary paths taken by *Penicillium glabrum* and *Botryosphaeria dothidea*, the diversity of Fusarium species, and the special evolutionary history of *Mucor racemosus*. Taken together, the phylogenetic results depict the potential of using combined full SSU and ITS sequences to distinguish between endophytic fungal species belonging to different genera. The level of the similarity of the obtained sequences ranged from 99-100%

**Fig 1.**
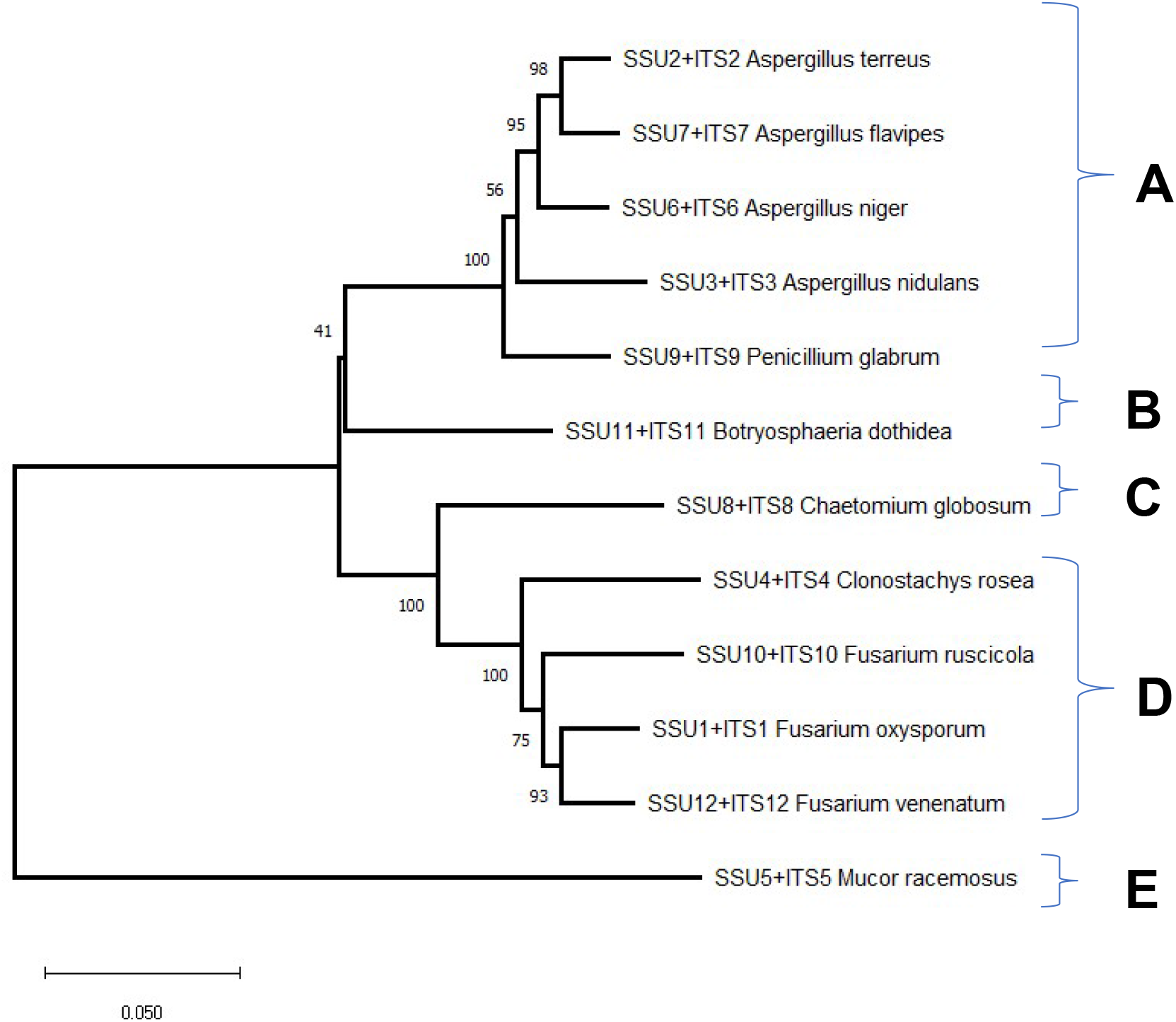
molecular phylogenetic analysis of ITS and SSU concatenated sequences of the isolated endophytic fungi from grapevine. The analysis was performed with MEGA using neighbor-joining method.

### 3.3 Bioagent control

#### 3.3.1. Inhibition of pathogens growth using bioagents endophytes by dual culture method

The data shown in Table 3 demonstrate the efficacy of the endophyte *P. maximus* as a bioagent in inhibiting the growth of *Ilyonectria destructans*,using which achieves a 70.83% inhibition rate. Antagonists’ rapid development is an important advantage in their competition for space, nutrition, and control over their prey hosts [24,25]. *A. alternative* had the lowest effect (7.87%) compared to other bioagents.

**Table 3.**
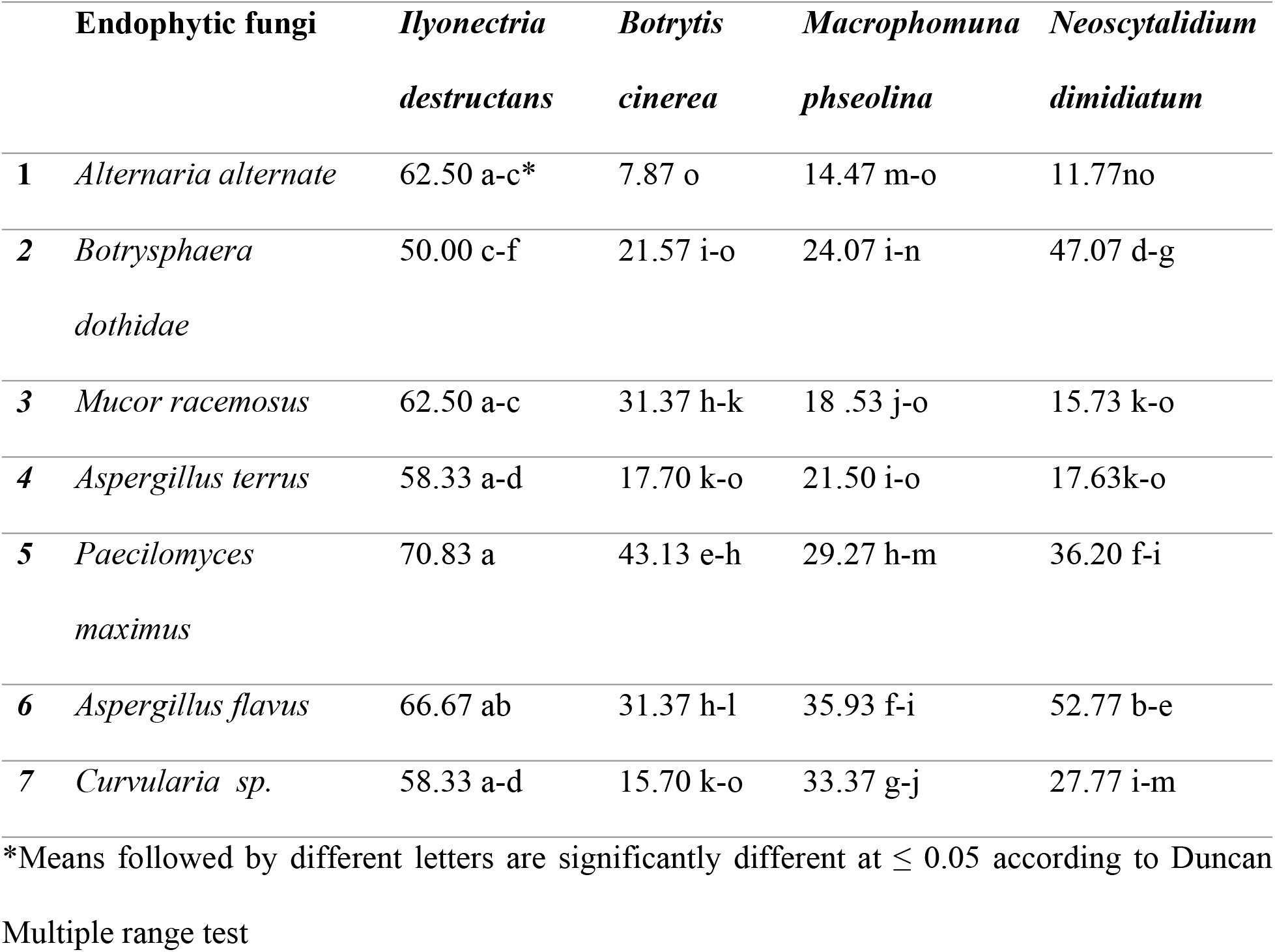
Inhibition percentage of pathogenic fungi using Dual culture method.

Figure (2) shows that *Aspergillus flavus* had the highest bioagent effect at 46.68%, primarily through competition for nutrients and space. *Paecilomyces maximus* came in second with 44.83%, while *Botryosphaeria dothidea* ranked third, inhibiting pathogen growth by 35.68%.

**Fig 2.**
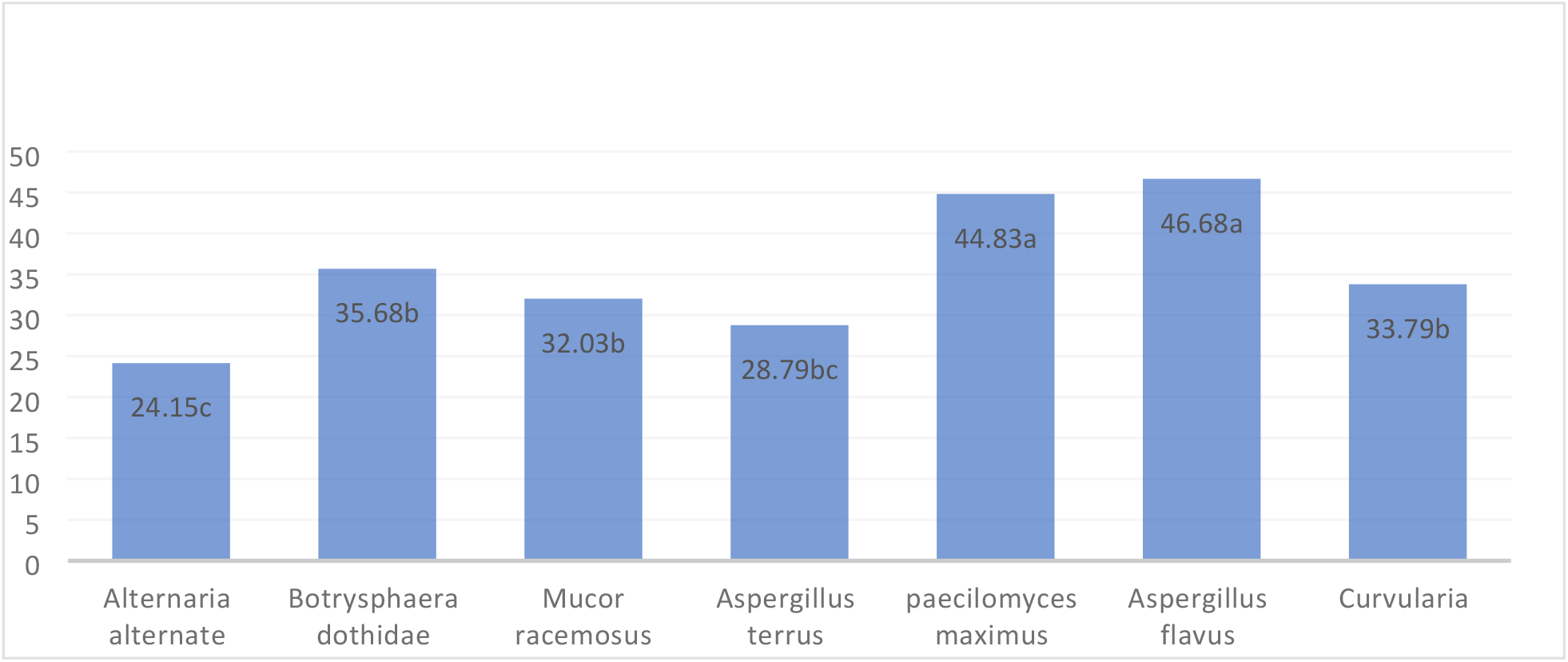
Inhibition percentage of pathogenic fungi growth in dual culture method.

#### 3.3.2 Food poisoning method

Table 4 Demonstrates that the endophytes *A. flavus* and *P. maximus* were the most effective bioagents in inhibiting the growth of *Ilyonectria destructans*, while *A. terrus, M. racemosus*, and *P. maximus* highly affected Botrytis cinerea’s growth (100% inhibition). Additionally, *M. racemosus* was found to be the most effective bioagent in inhibiting the growth of *Neoscytalidium dimidiatum* Table 4.

**Table 4.**
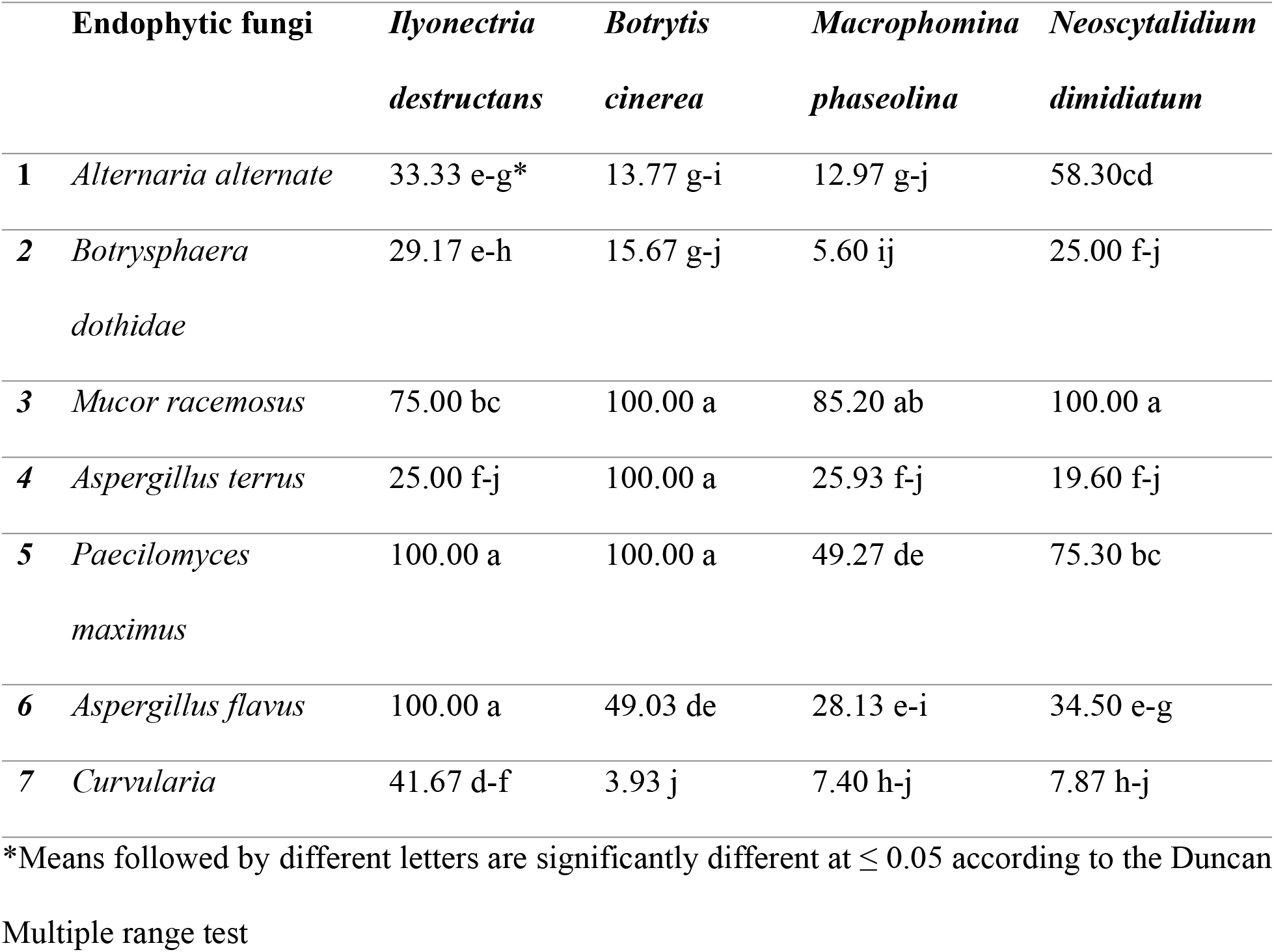
Mycelial growth diameter (mm) of pathogenic fungi treated with bioagents filtrate.

*Ilyonectria destructans* showed the greatest growth but the least resistance to most antagonists, while *Botrytis cinerea* was the most affected by the antagonistic process.

ITS and SSU primers used in the current investigation for molecular identification of endophytic fungi from grapevine at Duhok city-Iraq, screened in this study belonging genus Zygomycota was *Mucor racemosus*, with the highest occurrence in Besfki (100%) and the rest belong to Ascomycota, a most frequent endophytic fungi that comprise (*Aspergillus terreus, Aspergillus ustus, Aspergillus niger, Aspergillus flavipes, Botryosphaeria dothidea, Fusarium oxysporum, Fusarium ruscicol, Fusarium venenatum, Chaetomium globosum, Clonostachys rosea*, and *Penicillium glabrum*). Other endophytic species such as (*Acremonium,Alternaria, Arthrinium, Ascorhizoctonia, Aspergillus, Aureobasidium, Bipolaris, Botryosphaeria, Botrytis, Chaetomium,Cladosporium, Curvularia, Hypoxylon, Lasiodiplodia, Mycosphaerella, Nigrospora, Penicillium, Phoma, Scopulariopsis)* have already been identifying as endophytic fungi in the earlier study that isolated from stem of grapevine, in Beijing of China [26]. Endophytic fungi have been great potential as biocontrol agents against the diseases of grapevine that caused by pathogens. Isolating and identifying endophytes from a grapevine, evaluating their antagonistic activity against major pathogens, and investigating the mechanism behind their biocontrol potential. It emphasizes the importance of early detection of pathogen in vine yard nurseries that could help to prevent spreading of pathogen.

## 4. Conclusions

This study reveals the diversity and effectiveness of endophytic fungi from Vitis vinifera in the Duhok province-Iraq as biocontrol agents against grapevine diseases. Twelve fungal species were identified, with *Paecilomyces maximus, Aspergillus flavus*, and *Mucor racemosus* showing strong inhibition of major grapevine pathogens. These findings support the use of endophytic fungi as sustainable alternatives for disease management in vineyards, encouraging further research and practical applications in integrated pest management.

